# Cost Effective, Experimentally Robust Differential Expression Analysis for Human/Mammalian, Pathogen, and Dual-Species Transcriptomics

**DOI:** 10.1101/337527

**Authors:** Amol Shetty, Anup Mahurkar, Scott Filler, Claire M. Fraser, David A. Rasko, Vincent M Bruno, Julie C. Dunning Hotopp

## Abstract

As sequencing read length has increased, researchers have quickly adopted longer reads for their experiments. Here, we examine host-pathogen interaction studies to assess if using longer reads is warranted. Six diverse datasets encountered in studies of host-pathogen interactions were used to assess what genomic attributes might affect the outcome of differential gene expression analysis including: gene density, operons, gene length, number of introns/exons, and intron length. Principal components analysis, hierarchical clustering with bootstrap support, and regression analyses of pairwise comparisons were undertaken on the same reads, looking at all combinations of paired and unpaired reads trimmed to 36,54,72, and 101-bp. For *E coli,* 36-bp single end reads performed as well as any other read length and as well as paired end reads. For all other comparisons, 54-bp and 72-bp reads were typically equivalent and different from 36-bp and 101-bp reads. Read pairing improved the outcome in several, but not all, comparisons in no discernable pattern, such that using paired reads is recommended in most scenarios. No specific genome attribute appeared to influence the data. However, experiments with an *a priori* expected greater biological complexity had more variable results with all read lengths relative to those with decreased complexity. When combined with cost, 54-bp paired end reads provided the most robust, internally reproducible results across all comparisons. However, using 36-bp single end reads may be desirable for bacterial samples, although possibly only if the transcriptional response is expected *a priori* to be robust.

**DATA SUMMARY:**

1. The human only CSHL Encode data set (1) was downloaded from ftp://hgdownload.cse.ucsc.edu/goldenPath/hg19/encodeDCC/wgEncodeCshlLongRnaSeq/.
2. The data from mice vaginas infected with *Candida albicans* (2) was downloaded from the SRA (url - https://trace.ncbi.nlm.nih.gov/Traces/sra/?study=SRP057050).
3. The data from *Aspergillus fumigatus* cells in contact with human cells was downloaded from the SRA (url - https://www.ncbi.nlm.nih.gov/bioproject/399754).
4. The data from a strand-specific library from a study comparing *C. albicans* cells in contact with human cells with those in media (3) was downloaded from the SRA (url - https://trace.ncbi.nlm.nih.gov/Traces/sra/?study=SRP011085).
5. The data from *C. albicans* in culture media (3) was downloaded from the SRA (url - https://trace.ncbi.nlm.nih.gov/Traces/sra/?study=SRP011085).
6. The data from *Escherichia coli* grown in different media (4) was downloaded from the SRA (url - https://trace.ncbi.nlm.nih.gov/Traces/sra/?study=SRP056578).

I/We confirm all supporting data, code and protocols have been provided within the article or through supplementary data files. ⊠

**IMPACT STATEMENT:** As sequencing technologies improve, sequencing costs decrease and read lengths increase. We examine host-pathogen interaction studies to assess if using these longer reads is warranted given their increased cost relative to using the same number of shorter reads. To this end we compared the use of various read lengths and read pairing for six diverse host-pathogen datasets with varying genomic attributes including: gene density, operons, gene length, number of introns/exons, and intron length. We find that in the bacterial sample, 36-bp single end reads performed as well as any other read length and as well as paired end reads. When combined with cost, 54-bp paired end reads provided the most robust, internally reproducible results for all other comparisons. Read pairing improved the outcome in several, but not all, comparisons in no discernable pattern, such that using paired reads is recommended in most scenarios. No specific genome attribute appeared to influence the data.

## INTRODUCTION

As sequencing throughput has increased and sequencing costs have decreased, measuring differential expression of genes using sequence data has become an increasingly powerful, effective, and popular approach. While there are several derivations, particularly for the downstream analyses, essentially, a randomly sheared sequencing library is constructed from cDNA synthesized from the RNA samples of interest. Following the sequencing of millions of reads from these libraries, the transcript abundance is measured by counting the reads or sequencing depth underlying each transcript. A normalized version of this number that accounts for numerous factors including the gene length, total number of reads sequenced, and/or total number of reads mapping is then used to compare the samples of interest and to identify genes that are differentially expressed.

The most common platform used today for such analyses is the Illumina HiSeq, which currently generates ∼90 Gbp of 150-bp paired end reads for ∼$3000. This platform sees frequent updates yielding longer reads and decreasing costs per base pair (bp). As read lengths have increased, many researchers have quickly used the increased read lengths assuming it can only result in better data. However, despite decreasing costs per bp, ultimately the longer reads often mean an increased cost per read and typically fewer reads are sequenced for the same cost. While this leads to the same sequencing depth, it results in fewer independent measurements at each position. For example, a shift from 100-bp paired end reads to 125-bp paired end reads can lead to a 20% reduction in the number of reads sequenced to obtain the same sequencing depth. However, the decreased number of reads can actually result in reduced statistical power since a single read will contribute to the sequencing depth at a larger number of positions. Therefore, biases in the underlying reads may be amplified with longer reads.

One alternate approach is to sequence the same number of overall base pairs, but use shorter paired reads. Such an approach would yield more sequence reads underlying each transcript and therefore more independent measurements at each position. For example, the use of 50-bp paired end reads as opposed to 100-bp paired end reads would lead to a 200% increase in the number of reads sequenced to obtain the same sequencing depth. Another alternative would be to sequence single reads, as opposed to paired reads. However, both read length and read pairing are expected to influence the accuracy of read mapping, which is the crucial first step in any RNASeq analysis pipeline. Furthermore, these factors may influence various genomes differently. For example, paired reads may be more beneficial in a genome with a large number of para logo us genes, gene families, and or repeats.

Recently this was examined through an analysis of read lengths of various length and pairing status (paired v. unpaired) for a human transcriptome dataset that concluded that 50 bp single end reads could be used reliably for differential expression analysis, but that splice detection required longer, paired reads (1). However, what works best in human datasets may not always be best for other organisms. Therefore, and given the caveats described above, we sought to investigate the influence of read length and read pairing on differential expression analysis across a variety of genomes of various complexity including (a) genome size, (b) presence/absence of introns, (c) length of introns, (d) number of introns per gene, (e) number of genes, and (f) percentage of genes transcribed (**Table 1**). In several instances, we have increased the complexity to include sequencing data that contains both a mammalian host and an associated pathogen. Ultimately, the goal is to identify the most appropriate and most cost-effective sequencing strategy based on the intrinsic properties of the genome(s) being analyze. In this way, the available resources can be appropriately distributed in order to maximize the number of biological replicates for the conditions being examined while maintaining the greatest quality results.

**Table 1.** Data set attributes

| No. | Host | Pathogen | Mapping Target | Genome Size of Target | Phylogenetic Domain of Target | Median Intron Length in Targets | Genes in Target | Exons/Gene in Target |
| --- | --- | --- | --- | --- | --- | --- | --- | --- |
| 1 | Human | None | Human | 3.09 Gbp | Eukaryote | 1501 bp | 60,107 | ~5.4 |
| 2 | Mouse | Candida | Mouse | 2.73 Gbp | Eukaryote | 1286 bp | 43,346 | ~6 |
| 3 | Human | Aspergillus | Aspergillus | 29.4 Mbp | Eukaryote | 60 bp | 9,898 | ~2.9 |
| 4 | Human | Candida | Candida | 14.3 Mbp | Eukaryote | 87 bp | 8,254 | ~1.1 |
| 5 | None | Candida | Candida | 14.3 Mbp | Eukaryote | 87 bp | 8,254 | ~1.1 |
| 6 | None | E. coli | E. coli | 4.97 Mbp | Prokaryote | NA * | 4,917 | NA |

\*NA=not applicable

## METHODS

### Reference genomes

The human, mouse, and *Aspergillus* reference genomes (**Table 1**) in FASTA format and annotation files in GRF or GFF formats were downloaded from Ensembl, while the C. *albicans* and *E. coli* ones were downloaded from the *Candida* Genome Database (http://www.candidagenome.org) and NCBI, respectively. The FASTA genomic sequences were indexed using SAMTOOLS (v. 0.1.19) (5). The GTF/GFF reference annotations were used to extract genomic coordinates for the genes, exons and introns using the BEDTOOLS (v. 2.17.0) (6).

### Sequencing Data Used

The 101-bp paired end sequencing reads for each sample (Table 2) were trimmed from the 3 ‘end of the sequence read to generate 36-bp, 54-bp, and 72-bp reads using the FASTX-Toolkit (http://hannonlab.cshl.edu/fastx_toolkit) generating 2 separate FASTQ files consisting of first-in-pair reads and second-in-pair reads that were compressed for downstream analysis.

**Table 2.**
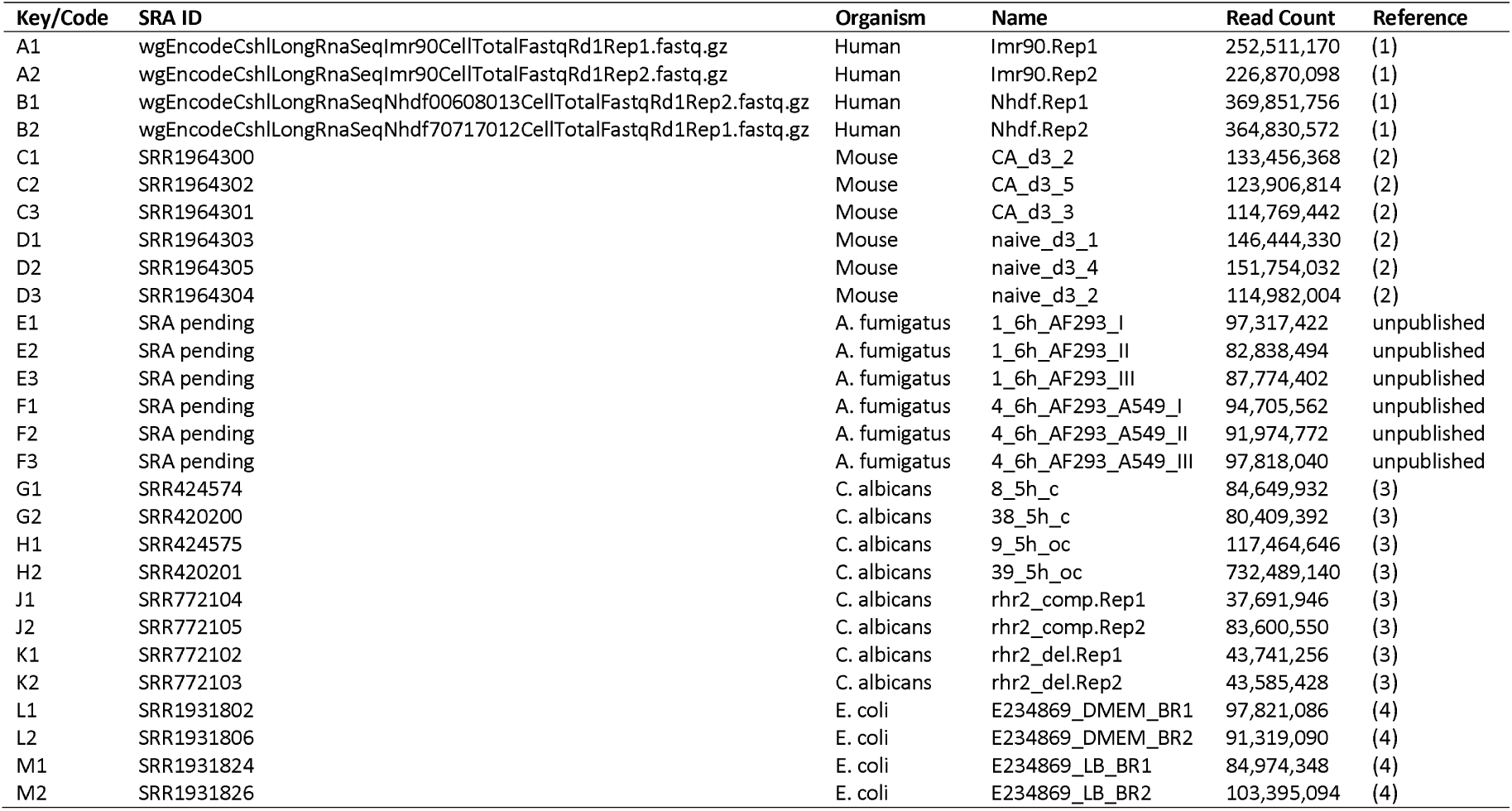
Sample key for comparisons in Differential Expression Analysis

### Reference based alignment

The sequencing reads were aligned to their respective reference genome FASTA sequence using the TopHat splice-aware aligner (v. 2.012) (7) for eukaryotic data or Bowtie aligner (v. 0.12.9) (8) for prokaryotic data allowing for a maximum of 2 mismatches per aligned read, an inner mate distance of 200 bp, and discarding reads that aligned to more than 20 genomic loci. The alignment files were sorted, indexed, and converted between BAM and SAM formats using SAMTOOLS (v. 0.1.19) (5). The alignment files were used to compute the total number of reads per sample, the number of reads that aligned to the reference genome, the number of reads that mapped once to the genome and the number of reads that mapped to >1 but <20 genomic loci (**Table 2**). The percentage of reads that mapped to exons, introns, genes and intergenic regions of the genome were computed based on coordinates from the respective annotation files in GTF/GFF format.

### RPKM calculations

The number of reads that mapped to each gene was calculated from the BAM alignments using HTSeq (v. 0.5.4) (9) and were further normalized for sequencing library depth and gene length to estimate the read counts per kilobase of the gene length per million mapped reads (RPKM) for each gene for each set of FASTQ files.

### Hierarchical clustering and PCA

The raw counts from HTSeq were further normalized using DESeq (v. 1.10.1) (10) in R (v. 2.15.2) (11). Genes with low read counts across all samples fora dataset were excluded from downstream analysis. The final set of normalized gene expression values for each gene for each sample within a dataset are used to compute a Euclidean distance matrix between every pair of samples that was used to generate a heat map cluster with PVCLUST with 1000 bootstraps. Eigen vectors were calculated with the PCA package in R to determine the first and second PCs that illustrate the vectors with the largest variance in the dataset.

The final set of normalized gene expression values for each gene for each sample within a dataset are used to test for differential gene expression between the two conditions using the ‘negative binomial’ test incorporated within DESeq (v. 1.10.1) (10) in R (v. 2.15.2) (11)). The final results have then been filtered to determine significant differentially expressed genes using a <5% false discovery rate (FDR), >2-fold-change, and a >10^th^ percentile of average normalized gene expression distribution within the dataset.

## RESULTS

### Design and Data Set Selection

We examined RNASeq data from six studies to test the effect of read length and read pairing on gene expression data from a wide set of host-pathogen samples, including (1) the human only CSHL Encode data set used in a prior analysis of the effect of read length on transcriptome analysis (1), (2) data from mice vaginas infected with *Candida albicans* (2), (3) unpublished data from a study comparing *Aspergillus fumigatus* cells in contact with human cells with those in media, (4) data from a strand-specific library from a study comparing C. *albicans* cells in contact with human cells with those in media (3), (5) data from *C. albicans* in culture media (3), and (6) data from *Escherichia coli* grown in different media (4) **(Table 1).** This includes eukaryotic and prokaryotic genomes; organisms of varying genome size and varying numbers of genes; organisms with and without introns; organisms of varying intron length; organisms with varying number of exons/gene; and data from single organisms compared to those from mixtures of organisms with an emphasis on host-pathogen systems **(Table 1).** All of the data sets used were generated as 101-bp paired end reads. Data was trimmed from the 30-end of the read to generate 36-bp, 54-bp, and 72-bp data sets for comparison. The first read in the pair was analyzed separately from the second read in the pair when single end reads were analyzed.

To examine the influence of read length and pairing at many steps, analyses were undertaken on multiple data sets. Mapping statistics were calculated from the Bowtie alignments. Principal components analysis (PCA) and hierarchical clustering were undertaken on FPKM values for each individual replicate in each biological condition **(Additional Files 1-12).** Scatterplots were used to examine differential expression results obtained with DESeq **(Additional Files 13-14)**.

### Read mapping as a function of read length

The number of reads mapping is dependent upon the number of mismatches allowed, as well as the uniqueness of the sequence, both of which are expected to vary by read length and the aligner used. In this case, Bowtie was used as the aligner, as it is the most prevalent aligner used for transcriptome studies today. With Bowtie, we expect that the number of the reads that map to multiple sites (multimap reads) will decrease with read length while the number of mismatches will increase with the read length. Therefore, we expect that fewer 36-bp reads will map uniquely since a greater proportion will multi-map, and we expect that fewer 101-bp reads will map because of the accumulation of sequencing errors, which increases with read length.

As expected, in half of the cases fewer reads map uniquely for 36-bp and 101-bp for both paired and single end reads, relative to the 54-bp and 72-bp equivalents **(Figure 1ABC).** The number of multi-map reads that do not map uniquely decreases as a function of read length **(Figure 1AB,** squares). However, in organisms with smaller genomes that have no introns (i.e. *E coii)* ora limited number of introns (i.e. *C. albicans*), increasing read length leads to decreasing mapped read counts **(Figure 1DEF)**.

**Figure 1.**
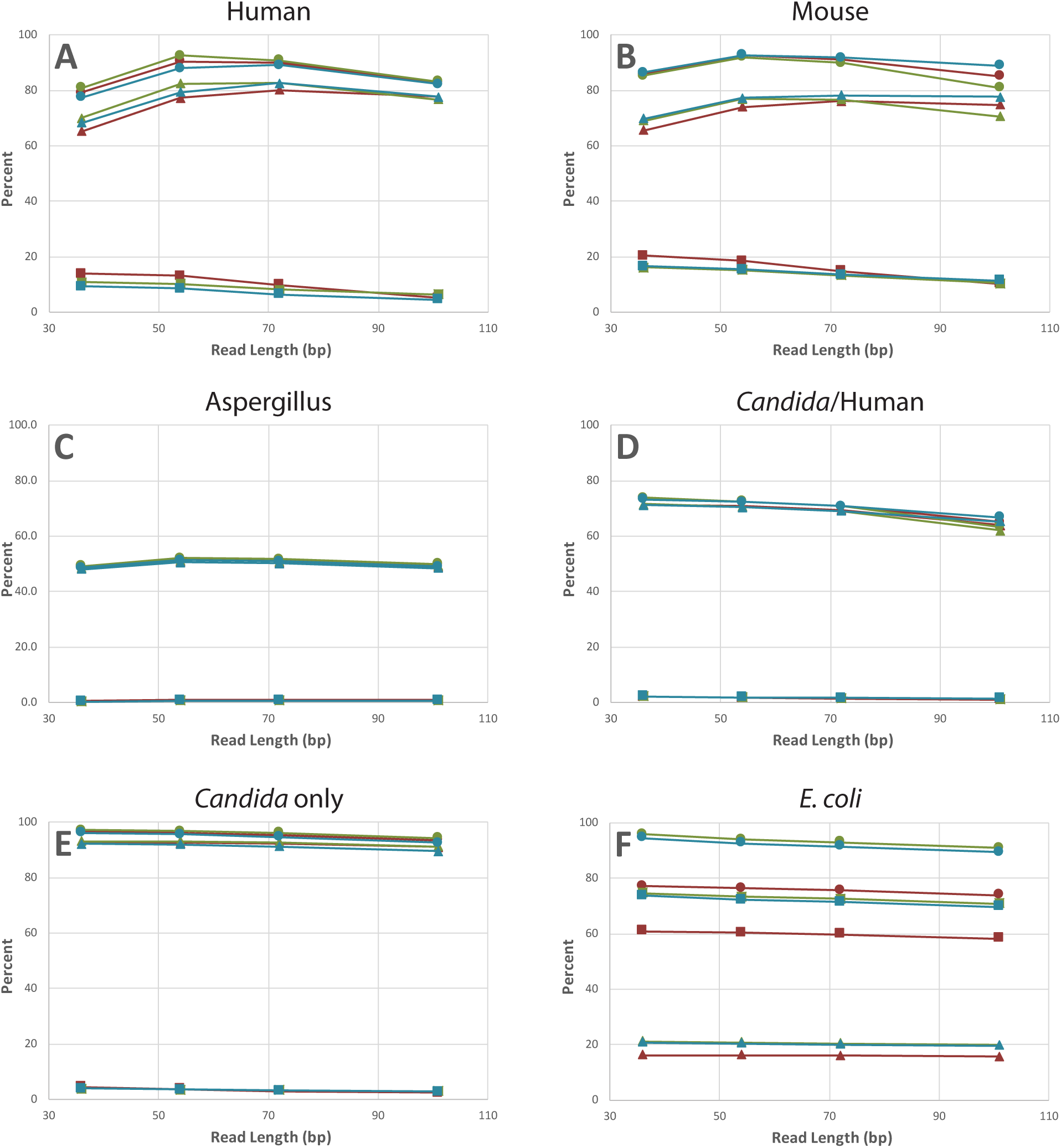
The average percentage of reads mapping (circles, left axis), reads mapping uniquely (triangles, left axis), and reads not mapping uniquely (squares, right axis) are compared for 36-bp, 54-bp, 72-bp, and 100-bp reads for the human (panel A), mouse (panel B), *Aspergillus* (panel C), *Candida*/host (panel D), *Candida* only (panel E), and *E coli* (panel F) data sets. Results are compared for mappings with the paired reads (red), only the first read in the pair (green), and only the second read in the pair (blue).

The greatest proportion of multi-mapping reads were found in *E coli* followed by mouse and human. Unlike the eukaryotic datasets analyzed where polyadenylated RNA can be enriched and sequenced, the *E coli* data had a sizable proportion of rRNA left that was sequenced. Given that there are 7 copies of the rRNA in the reference genome used for mapping (12), a large number of multi-mapping reads were expected. Therefore, as expected, >99% of reads mapping to the rRNA genes were multi-mapping reads, and on average, 78% of the mapped reads mapped to the rRNA genes. The increase in multi-mapping reads in human and mouse is expected given their genome size and composition. In both humans and mice, the paired end reads yielded slightly more multiple hits than the single end reads, which we attribute to how the aligner handles multi-mapping reads.

### PC analysis of read length

If read length is of no consequence, samples of the various read lengths should be more similar to one another than to samples from other biological conditions or replicates, which can be examined with PCA. In that case, we would expect the first principle component (PC) to separate the data based on biological condition and the second PC to separate the data based on replicates. Furthermore, we would expect all of the read lengths derived from the same data to be tightly grouped. This was observed for *E coli* paired end reads **(Figure 2A).** It was also observed for the for the other *E coli* comparisons **(Additional File 6),** *Candida* paired end reads **(Additional File 4),** and to a lesser degree the *Aspergillus* comparisons **(Additional File 3)**.

**Figure 2.**
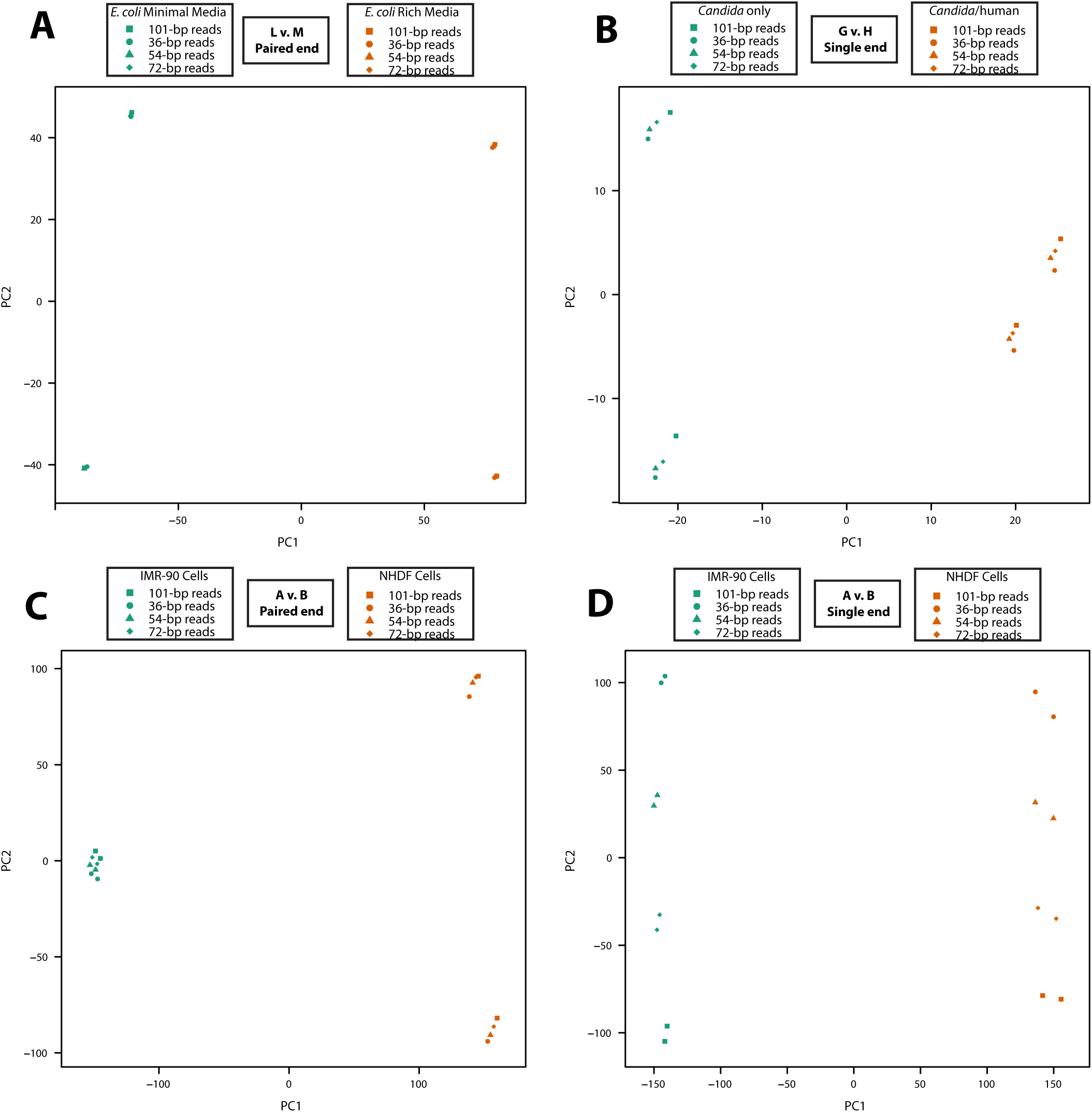
A PCA was undertaken for a vector representing data for the different read lengths (circle, 36-cp; triangle, 54-bp; diamond, 72-bp; square, 101-bp), replicates (green v. red), and biological conditions. Four representative results are illustrated here with *E coli* paired end data (panel A), *Candida*/human first-in-read single end reads (panel B), human paired end reads (panel C), and human first-in-read single end reads (panel D). All PCA plots for read length are provided in **Additional Files 1-6** and pairing status are provided in **Additional Files 7-12.**

However, in some cases the read length played a greater role. The single end reads from the *Candida*/human data set demonstrate similar PC1 and PC2, but the spread of the data points suggests that read length may have some influence on the data **(Figure 2B).** This was also observed for the paired end reads and single end reads for the other *Candida* data set **(Additional File 5)** as well as the paired end reads from the human/CSHL data set **(Additional File 1)**.

The influence of read length is very pronounced in the single end reads from the CSHL data set, which were separated on the first PC by biological replicate, but were separated by read length on the second PC **(Figure 2D, Additional File 1).** This suggests that there were greater distinctions in the length of the read pairs than there were in the replicates. This was also true for both the paired end reads and single end reads from the *Candida*-infected mouse vagina data set **(Additional File 2).** When read length does divide the data, it is distinguished from decreasing to increasing read length along the axis, as opposed to a random order.

### Hierarchical clustering as a function of read length

In numerous cases, hierarchical clustering (complete clustering, correlation distance) of the datasets with statistical support (AU, approximately unbiased and BP, bootstrap probability) is consistent with the PCA. When the PCA reveals data clustering by biological condition and then replication, but not by read length, in the *E coli* datasets, the heat map and dendrogram show similar, well-supported (confidence ≥80%) clustering **(Figure 3A, Additional File** 6). And in the instances where the PCA analysis revealed that read length had the greatest influence, the hierarchical clustering showed the greatest variability in clustering. This was most striking with the mouse data, which had poor clustering of the data, with no discernable pattern **(Figure 3B, Additional File 2).** While the mouse samples clustered by condition, in many instances data with the same read length but from different replicates clustered better than data from the same replicates with different read lengths. This suggests that the read length is influencing the data. However, between these extremes the hierarchical clustering showed more granularity and in most cases some clustering by read length instead of replicates was found **(Additional Files 1, 2,4, 5),** particularly for the 36 bp reads.

**Figure 3.**
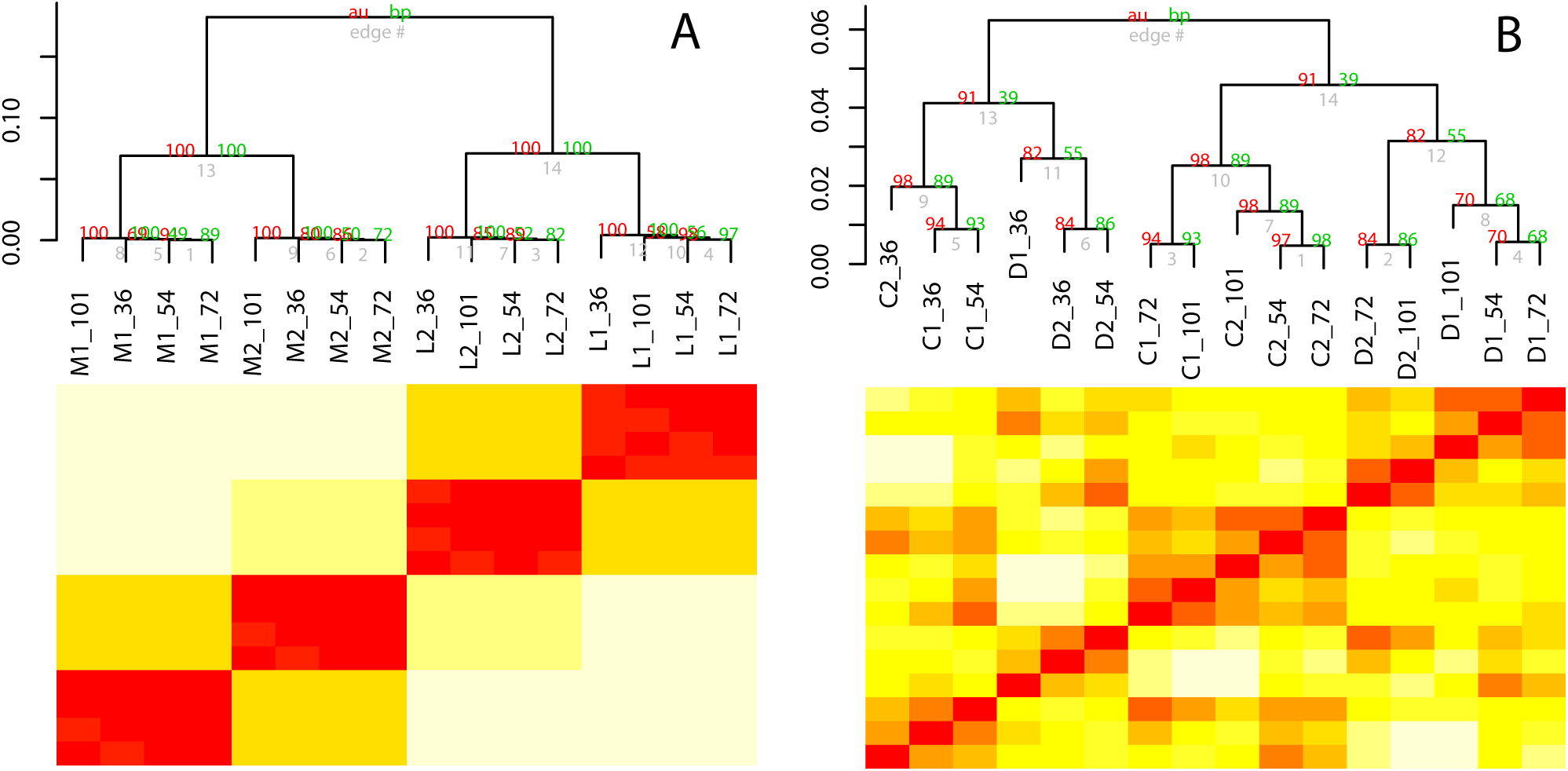
Hierarchical clustering using PVCIustfor bootstrap support was undertaken for a vector representing data for each sample at different read lengths. Samples are labeled according to the key in Table 2 followed by the read length (36-bp, 54-bp, 72-bp, and 101-bp). Two representative results are illustrated here with (A) *E coli* and (B) mouse paired end data. In the *E coli* data, read length did not affect the clustering of the data, while the largest effect of read length was observed with the mouse data.

### Log-fold Change of differentially expressed genes as a function of read length

For all comparisons, the log-fold change of differentially expressed genes between the two conditions correlates well between two replicates with R^2^ values ranging from 0.63 to 1.0 (average: 0.92; median: 0.95) across all pairwise comparisons of read length for single end and paired end reads **(Table 3)**.

**Table 3.** R^2^ Values for All Pairwise Comparisons of Read Length

| Experiment | Pairing Status | 36 v. 54 | 36 v. 72 | 36 v. 101 | 54 v. 72 | 54 v. 101 | 72 v. 101 |
| --- | --- | --- | --- | --- | --- | --- | --- |
| A v. B | paired | 0.95 | 0.92 | 0.88 | 0.97 | 0.93 | 0.95 |
| A v. B | single 1 | 0.94 | 0.91 | 0.88 | 0.95 | 0.91 | 0.95 |
| A v. B | single 2 | 0.95 | 0.93 | 0.89 | 0.96 | 0.93 | 0.95 |
| C v. D | paired | 0.89 | 0.93 | 0.93 | 0.94 | 0.93 | 0.97 |
| C v. D | single 1 | 0.75 | 0.73 | 0.72 | 0.98 | 0.96 | 0.98 |
| C v. D | single 2 | 0.95 | 0.97 | 0.96 | 0.92 | 0.9 | 0.99 |
| F v. E | paired | 0.87 | 0.87 | 0.84 | 1 | 1 | 0.97 |
| F v. E | single 1 | 0.94 | 0.93 | 0.93 | 1 | 1 | 1 |
| F v. E | single 2 | 0.94 | 0.85 | 0.93 | 0.99 | 0.99 | 1 |
| H v. G | paired | 0.9 | 0.8 | 0.84 | 0.88 | 0.82 | 0.92 |
| H v. G | single 1 | 0.84 | 0.64 | 0.63 | 0.79 | 0.66 | 0.78 |
| H v. G | single 2 | 0.9 | 0.75 | 0.67 | 0.89 | 0.77 | 0.8 |
| K v. J | paired | 1 | 0.98 | 0.99 | 0.99 | 1 | 1 |
| K v. J | single 1 | 0.99 | 0.98 | 0.98 | 0.98 | 0.99 | 0.99 |
| K v. J | single 2 | 0.86 | 0.77 | 0.96 | 0.93 | 0.93 | 1 |
| L v. M | paired | 1 | 1 | 1 | 1 | 1 | 1 |
| L v. M | single 1 | 1 | 1 | 0.99 | 1 | 0.99 | 1 |
| L v. M | single 2 | 1 | 0.99 | 0.99 | 0.99 | 0.99 | 1 |

Remarkably, all such pairwise comparisons with *E coli* yield R^2^ values of 0.99 or 1.00 **(Figure 4),** suggesting that 36-bp single end reads yield the same results as 101-bp paired end reads. Overall though, comparisons that include 36-bp reads are typically not as good as those with the long read lengths **(Table 3).** Of the remaining comparisons, the best correlations are found in comparisons of the closest read lengths, specifically 54 bp v. 72 bp and 72 bp v. 101 bp) **(Table 3)**.

**Figure 4.**
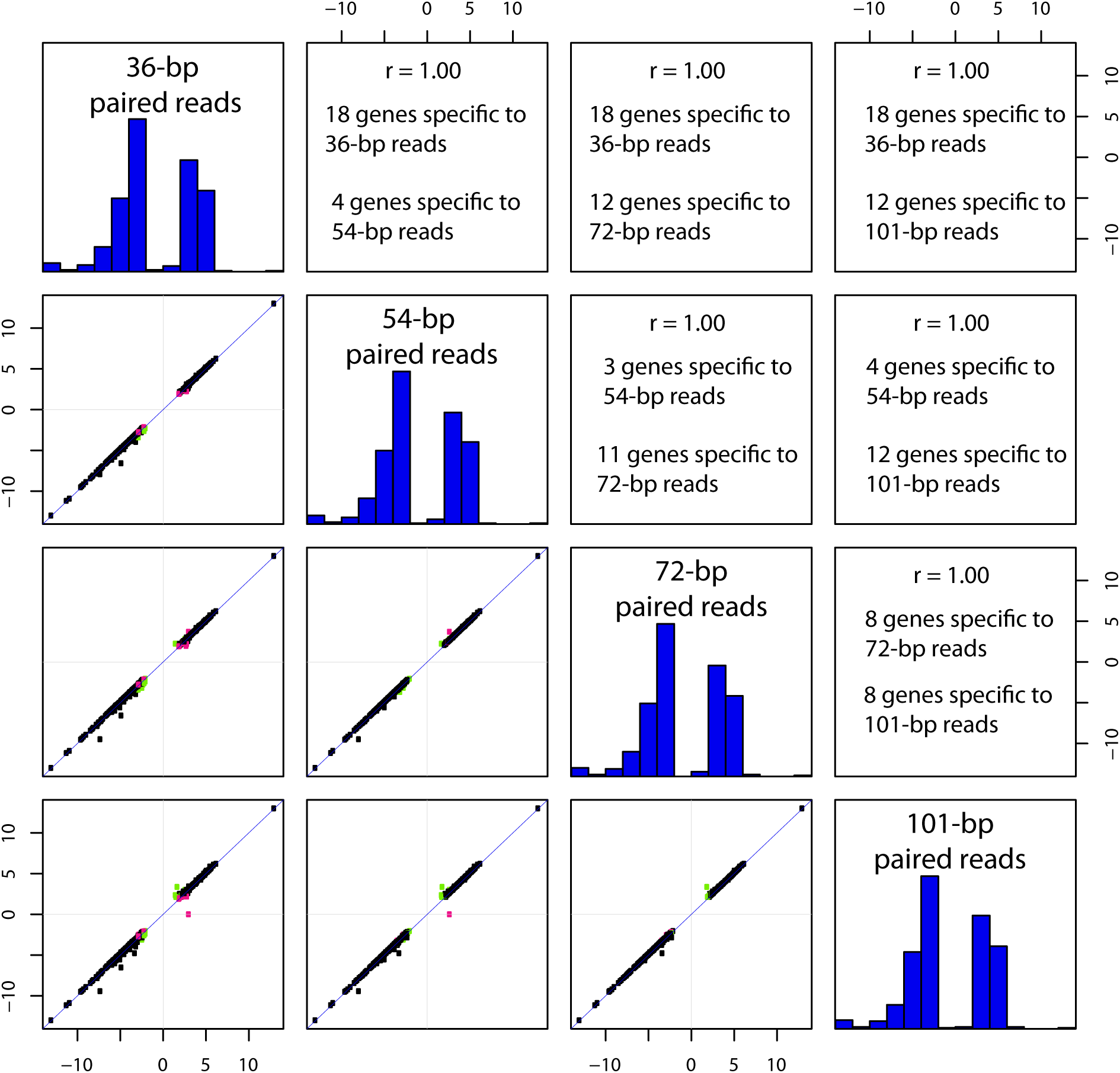
The differentially expressed genes identified in *E coli* **(L** v. **M)** using an adjusted p-value (FDR) cutoff ≤0.05 for paired end reads at varying read lengths within a dataset were compared using Pearson’s correlation implemented in the R statistical tool and illustrated as a matrix of scatterplots. The diagonal represents the histogram of log-transformed fold-changes within the comparison. The lower plots represent the correlation between comparisons with singleton DEGs identified forcompariosns on the x-axis (pink) and y-axis (green). Genes with FDR > 0.05 in both comparisons are not shown. The upper portion of the plot lists the corresponding Pearson’s correlation coefficient and the number of singleton DEGs identified in each comparison.

A slightly different result is observed when focusing on genes found to be differentially regulated at one read length but not found to be differentially regulated at another read length, referred to as singletons. In this case, the 54-bp v. 72-bp comparison consistently outperformed all other comparisons **(Table 4).** The next best comparisons were the other two groupings of similar sizes, 36-bp v. 54-bp and 72-bp v. 101-bp **(Table 4).**

**Table 4.** Number of Singletons for All Pairwise Comparisons of Read Length

| Experiment | Pairing Status | 36 v. 54 | 36 v. 72 | 36 v. 101 | 54 v. 72 | 54 v. 101 | 72 v. 101 |
| --- | --- | --- | --- | --- | --- | --- | --- |
| A v. B | paired | 1540 | 2010 | 2704 | 1060 | 2030 | 1478 |
| A v. B | single 1 | 1500 | 2088 | 3017 | 1248 | 2377 | 1705 |
| A v. B | single 2 | 1619 | 2173 | 2936 | 1162 | 2201 | 1591 |
| C v. D | paired | 59 | 60 | 74 | 47 | 59 | 50 |
| C v. D | single 1 | 164 | 170 | 164 | 44 | 56 | 48 |
| C v. D | single 2 | 73 | 72 | 85 | 49 | 68 | 43 |
| F v. E | paired | 90 | 16 | 18 | 7 | 7 | 8 |
| F v. E | single 1 | 16 | 15 | 14 | 5 | 8 | 7 |
| F v. E | single 2 | 19 | 16 | 15 | 9 | 8 | 5 |
| H v. G | paired | 131 | 169 | 268 | 84 | 193 | 149 |
| H v. G | single 1 | 207 | 294 | 445 | 167 | 344 | 295 |
| H v. G | single 2 | 205 | 301 | 428 | 154 | 329 | 243 |
| K v. J | paired | 13 | 19 | 23 | 12 | 14 | 10 |
| K v. J | single 1 | 17 | 23 | 29 | 12 | 20 | 24 |
| K v. J | single 2 | 19 | 23 | 38 | 6 | 25 | 25 |
| L v. M | paired | 22 | 30 | 30 | 14 | 16 | 16 |
| L v. M | single 1 | 37 | 46 | 69 | 19 | 52 | 47 |
| L v. M | single 2 | 36 | 57 | 74 | 41 | 56 | 47 |

### PC analysis of read pairing

Read pairing is expected to exert influences in many of the same ways as read length. Theoretically, a pair of 36-bp reads should provide benefits greater than a single 72-bp read, since 36-bp paired end reads will have 72-bp of specific sequence, as well as some information about the approximate distance between the two 36-bp reads. As such a pair of 36-bp reads could resolve repeats of a similar length to the insert size distribution of the library. If read pairing is of no consequence, paired and single end reads should be more similar to one another than to samples from other biological conditions or replicates, which can be examined with a PCA and hierarchical clustering, as was conducted for read length. For the PCA, each read length data set is examined separately for a given pairing status, and we would expect the first PC to separate the data based on biological condition and the second PC to separate the data based on replicates. This was observed for reads from the *E coli* data sets where read pairing did not matter for each of the four read lengths examined **(Additional File 12).**

In all other cases, at least some read lengths showed data being separated by pairing status instead of replication in the second PC. In the *Candida*-only data set, read pairing influenced the data for the 36-bp, 54-bp, and 72-bp paired end reads **(Additional File 11).** For Cand/da/human, read pairing influenced the data for the 36-bp paired end reads **(Additional File 10).** For the mouse vaginas, read pairing influenced the data for 36-bp and 54-bp reads with some effect seen with 72-bp reads **(Additional File 8).** And in the CSHL data set, some effect was seen at all read lengths **(Additional File 7).**

### Hierarchical clustering as a function of read pairing

Hierarchical clustering of the datasets largely supports the PCA analysis for read pairing. When the PCA reveals data clustering by biological condition and then replication, but not by read pairing, as is the case with the *E coli* datasets, the heat map and dendrogram show similar clustering, which is well supported by the AU/BP values (100%) **(Additional File 12).**

The instances where the PCA analysis showed the greatest influence of read pairing also showed the greatest variation in hierarchical clustering. In the 36-bp and 54-bp CSHL human reads, the samples were separated by biological condition with 100% support **(Additional File 7).** However, in one of the conditions, the reads were distinguished into three groups comprised of paired reads, first-in-pair reads, and second-in-pair reads, each with >90% support **(Additional File 7).** In the 72-bp reads, the node of the paired reads and the first-in-pair reads becomes poorly supported (<60% support) **(Additional File 7)**. However, unlike the PCA analysis, in the 101-bp reads, the samples are clustered by biological condition, then replication, and then read pairing, which might not be expected from the PCA analysis, where some influence of pairing was observed **(Additional File 7)**. This suggests the differences observed in the PCA analysis of pairing for the 101-bp reads can be resolved in the hierarchical clustering.

This difference between the PCA analysis and the hierarchical clustering is also seen in the mouse vagina dataset (**Additional File 8**). In the PCA analysis, the mouse vaginas showed a strong influence of read pairing in the 36-bp and 54-bp reads, as well as some influence of read pairing in the 72-bp reads (**Additional File 8**) that was similar to that seen in the CSHL dataset In the mouse vagina dataset, hierarchical clustering of the 72-bp and the 101-bp datasets resolved the biological conditions, then the replicates, and then the pairing status (**Additional File 8**).

Despite the differences in the PCA for the *Candida*/human dataset (**Additional File 10**) and the *Candida-only* dataset (**Additional File 11**), only the 101-bp reads were resolved first by biological condition, then replication, and then read pairing for both of these data sets. At all other read lengths, clusters separated by read pairing before replication (**Additional Files 10 & 11**).

## DISCUSSION

As sequencing technologies have improved, sequencing reads have become longer and the inclination is to use these longer sequencing reads to obtain presumably better data. However, with increasing read lengths also comes increasing costs. Here, we examine whether using longer reads provides a benefit when conducting differential expression transcriptomics experiments, or if the increased costs could be better spent in other ways, like increasing the number of reads sequenced or increasing sequencing of replicates. To this end, we compared six diverse datasets consisting of pairwise comparisons of two samples with at least two replicates per sample, frequently focusing on datasets encountered in studies of host-pathogen interactions. In all cases, the number of sequencing reads for each comparison remained constant between the comparisons, but the reads were trimmed to generate paired reads of four different lengths - 36 bp, 54 bp, 72 bp, and 101 bp.

For *E*. *coli,* 36-bp single end reads performed as well as any other read. Given the decreased cost, there does not seem to be any scientific justification for longer sequencing reads, or paired reads, for pairwise differential expression analyses based on this analysis. As read length increases, fewer reads map, likely owing to the known accumulation of errors in long reads. There are a sizable number of multi-mapping reads, due to the presence of rRNA in the comparison, but this does not appear to affect the results. In the PCA and hierarchical clustering, the data are always grouped first by biological condition, then technical replication, and then sequencing reads or pairing, demonstrating that read length and read pairing are less impactful than technical replication for this sample set. All pairwise comparisons of differentially expressed genes for read length or pairing yielded R^2^-values of 0.99 to 1.00. The only difference observed between read length or pairing is the number of singletons between results for each read length. Singletons are genes that found to be differentially regulated under one condition (in this case one read length) but not differentially regulated under another condition (in this case a different read length). Here we do observe a difference between the single end and paired end reads, with more singletons identified with the single end reads. However, these are largely genes that fall on the diagonal in pairwise plots, with differential expression levels that places them closer to the fold-threshold cutoff that they may fall over the threshold for the paired end reads and under the threshold in the single end reads, or vice versa. It is not possible to say which result is correct. In these cases, obtaining more sequencing reads and/or replicates may be more beneficial at resolving the significance of differential expression. As such, 36-bp single end reads seem to be the best, regardless of cost. Further work is needed to see if this result is widely applicable to other bacterial species and systems as well as more heterogenous populations of bacterial cells.

*Aspergillus* and *Candida*-only also yielded very similar results. The pairwise comparisons had very strong correlations, with typical R^2^ values of 0.99 or 1.00. The PCA and hierarchical clustering largely clustered by biological condition first, then replicate, and then read length or read pairing. In the hierarchical clustering there are instances where the technical replicate cluster together as opposed to with the biological condition, specifically the third replicates for the *Aspergillus* data. However, in these cases the clustering does not have strong statistical support. Collectively, more incongruences are observed with 36-bp and 101-bp reads than with 54-bp and 72-bp reads. As such some preference should be given to 54-bp and 72-bp reads, as this likely indicates that these read lengths yield the most robust results.

On the other end of the spectrum is the mouse transcriptome data. In this data, the number of multimapping reads decreased with increasing read length, showing an advantage to having longer reads. However, the total number of reads mapping decreased with increasing read length, likely owing to errors that accumulate in the reads that make them more difficult to map. The two middle read lengths (54-bp and 72-bp) performed best in terms of mapping percentage. For paired end and single end reads, in the PCA the data clusters (a) first by biological condition and then by read length or (b) first by biological condition and then by a mixture of read pairing and replication suggesting that the read length is strongly influencing the data. Hierarchical clustering of data separated by pairing status reveals that read length plays a larger role than even the biological condition with shorter reads clustering separately than longer reads, and within these clusters reads clustering by read length instead of replicate. Hierarchical clustering of data separated by read length reveals that at greater read lengths clustering is as expected --first by biological condition, then by replicate, and lastly by pairing status. However, at the two lower read lengths there was clustering first by biological conditions followed by a mixture of ^1^ clustering by pairing status as opposed to replicate. Analysis of the pairwise comparisons reveals that there is a particularly strong difference in the 36-bp first-in-pair single end reads with poorer R^2^ read lengths, and that this is due to genes having ratios over the ratio threshold in the 36-bp data but ratios near 1 in the 54-bp, 72-bp, or 101-bp data. In this case, clearly the 36-bp first-in-pair single end reads are yielding different results than all other comparisons, but what is the best data? While there are differences observed, we cannot assume that longer is necessarily better, as might be indicated with the decreasing mapping percentages as a function of read length. It might be important to consider this mouse transcriptome case unusual, possibly there was an undetected problem in the sequencing of the first read. But regardless the congruence between 54-bp, 72-bp, or 101-bp reads likely indicates that these read lengths yield the most robust results. Of these three there does not seem to be a read length that is clearly superior.

The remaining two comparisons (human cell lines only and *Candida* in differential contact with human cells) were both more variable than the other *Candida* sample, *E coli, or Aspergillus,* but without an obvious bias like the mouse transcriptome data. The samples always clustered first by biological condition, and then usually by replicate. However, 36-bp reads were sometimes found to cluster together rather than clustering by replicates as were 101-bp reads and paired end reads. The 54-bp and 72-bp reads were most likely to cluster as expected and were most similar to one another.

We intentionally chose to compare six data sets, representing a diverse array of genomic complexity to assess what attributes might affect the outcome. Our selection included genomes with high gene density, genomes with operons, genomes with long genes, genomes with many introns/exons, and genomes with long introns. We did not observe any obvious patterns associated with these criteria. *Aspergillus* has long introns, many introns/exons per gene, and a lower transcriptional density, yet it performed almost as well as *E coli* which has a greater transcriptional density, operons, and intronless genes. Instead the experiments roughly clustered into two groups that could be defined by the biological complexity of the transcriptional response. The group contained a comparison of a single synchronized culture growing on two different media and two comparisons of two cultures of mutants synchronized and growing on the same media. In these cases, the cultures were synchronized and as such the transcriptional response is expected to be well delineated. On the other hand, there was the transcriptional response of mice vaginal cells, unsynchronized human cell cultures, and *Candida* in the presence/absence of human cells. In these cases, the transcriptional response is likely to be less delineated and noisier, reflecting the lack of synchronicity of the cells and the increased diversity of the transcriptional response. It seems likely that in these cases, subtle changes that occur when altering the read length are altering the statistical significance of results as opposed to a drastic change in the measured fold-change of the response, which is largely observed in the pairwise comparisons. This suggest that in these cases, more sequencing reads, rather than longer sequencing reads, may allow for more robust conclusions to be drawn.

## Acknowledgements

We would like to thank other members of the IGS GCID project for their helpful suggestions, particularly other members of the Technology Core.

## Ethics approval and consent to participate

Not applicable.

## ABBREVIATIONS

Approximately unbiased (AU)

Bootstrap probability (BP)

Base pair (bp)

False Discovery Rate (FDR)

Principal component (PC)

Principal components analysis (PCA)

Read counts per kilobase of the gene length per million mapped reads (RPKM)

---

**Additional File 1. Compendium of figures for Encode CSHL comparisons of IMR-90 v. NHD cells with results separated by read pairing status.** A heatmap with hierarchical clustering with statistical support is shown on page 1 with the condition denoted according to letter code from Table 2, followed by the replicate designation and the read length. A PCA plot is shown on page 2 where the conditions are denoted by the shape (circle, IMR-90; triangle, NHD) and the read length by the color (green, 36 bp; blue, 54 bp; magenta, 72 bp; purple, 101 bp). On both pages, results are shown in the three panels for (A) paired end reads, (B) first-in-pair single end reads, and (C) second-in-pair single reads.

**Additional File 2. Compendium of figures for data from *Candida*-infected mouse vaginas with results separated by read pairing status.** A heatmap with hierarchical clustering with statistical support is shown on page 1 with the condition denoted according to letter code from Table 2, followed by the replicate designation and the read length. A PCA plot is shown on page 2 where the conditions are denoted by the shape (circle, CA_d3; triangle, naïve_d3) and the read length by the color (green, 36 bp; blue, 54 bp; magenta, 72 bp; purple, 101 bp). On both pages, results are shown in the three panels for (A) paired end reads, (B) first-in-pair single end reads, and (C) second-in-pair single reads.

**Additional File 3. Compendium of figures for *A. fumigatus* data with results separated by read pairing status.** A heatmap with hierarchical clustering with statistical support is shown on page 1 with the condition denoted according to letter code from Table 2, followed by the replicate designation and the read length. A PCA plot is shown on page 2 where the conditions are denoted by the shape (circle,1_6h_AF293; triangle, 4_6h_AF293) and the read length by the color (green, 36 bp; blue, 54 bp; magenta, 72 bp; purple, 101 bp). On both pages, results are shown in the three panels for (A) paired end reads, (B) first-in-pair single end reads, and (C) second-in-pair single reads.

**Additional File 4. Compendium of figures for *Candida-human* data with results separated by read pairing status.** A heatmap with hierarchical clustering with statistical support is shown on page 1 with the condition denoted according to letter code from Table 2, followed by the replicate designation and the read length. A PCA plot is shown on page 2 where the conditions are denoted by the shape (circle, 5h_c; triangle, 5h_oc) and the read length by the color (green, 36 bp; blue, 54 bp; magenta, 72 bp; purple, 101 bp). On both pages, results are shown in the three panels for (A) paired end reads, (B) first-in-pair single end reads, and (C) second-in-pair single reads.

**Additional File 5. Compendium of figures for *Candida-on\y* data with results separated by read pairing status.** A heatmap with hierarchical clustering with statistical support is shown on page 1 with the condition denoted according to letter code from Table 2, followed by the replicate designation and the read length. A PCA plot is shown on page 2 where the conditions are denoted by the shape (circle, rhr2_comp; triangle, rhr2_del) and the read length by the color (green, 36 bp; blue, 54 bp; magenta, 72 bp; purple, 101 bp). On both pages, results are shown in the three panels for (A) paired end reads, (B) first-in-pair single end reads, and (C) second-in-pair single reads.

**Additional File 6. Compendium of figures for *E. coli* data with results separated by read pairing status.** A heatmap with hierarchical clustering with statistical support is shown on page 1 with the condition denoted according to letter code from Table 2, followed by the replicate designation and the read ^1^ length. A PCA plot is shown on page 2 where the conditions are denoted by the shape (circle, DMEM; triangle, LB) and the read length by the color (green, 36 bp; blue, 54 bp; magenta, 72 bp; purple, 101 bp). On both pages, results are shown in the three panels for (A) paired end reads, (B) first-in-pair single end reads, and (C) second-in-pair single reads.

**Additional File 7. Compendium of figures for Encode CSHL comparisons of IMR-90 v. NHD cells with results separated by read length.** A heatmap with hierarchical clustering with statistical support is shown on page 1 with the condition denoted according to letter code from Table 2, followed by the replicate designation and the pairing status such that (0) paired reads, (1) first-in-read single end read, and (2) second-in-read single end read. A PCA plot is shown on page 2 where the conditions are denoted by the shape (circle, IMR-90; triangle, NHD) and the pairing status by the color (green, paired end; blue, first-in-pair single end read; magenta, second-in-pair single end read). On both pages, results are shown in the four panels: (A) 36-bp reads, (B) 54-bp reads, (C) 72-bp reads, and (D) 101-bp reads.

**Additional File 8. Compendium of figures for data from *Candida*-infected mouse vaginas with results separated by read length.** A heatmap with hierarchical clustering with statistical support is shown on page 1 with the condition denoted according to letter code from Table 2, followed by the replicate designation and the pairing status such that (0) paired reads, (1) first-in-read single end read, and (2) second-in-read single end read. A PCA plot is shown on page 2 where the conditions are denoted by the shape (circle, CA_d3; triangle, naïive_d3) and the pairing status by the color (green, paired end; blue, first-in-pair single end read; magenta, second-in-pair single end read). On both pages, results are shown in the four panels: (A) 36-bp reads, (B) 54-bp reads, (C) 72-bp reads, and (D) 101-bp reads.

**Additional File 9. Compendium of figures for *A. fumigatus* data with results separated by read length.** A heatmap with hierarchical clustering with statistical support is shown on page 1 with the condition denoted according to letter code from Table 2, followed by the replicate designation and the pairing status such that (0) paired reads, (1) first-in-read single end read, and (2) second-in-read single end read. A PCA plot is shown on page 2 where the conditions are denoted by the shape (circle, l_6h_AF293; triangle, 4_6h_AF293) and the pairing status by the color (green, paired end; blue, first-in-pair single end read; magenta, second-in-pair single end read). On both pages, results are shown in the four panels: (A) 36-bp reads, (B) 54-bp reads, (C) 72-bp reads, and (D) 101-bp reads.

**Additional File 10. Compendium of figures for *Candida-human* data with results separated by read length.** A heatmap with hierarchical clustering with statistical support is shown on page 1 with the condition denoted according to letter code from Table 2, followed by the replicate designation and the pairing status such that (0) paired reads, (1) first-in-read single end read, and (2) second-in-read single end read. A PCA plot is shown on page 2 where the conditions are denoted by the shape (circle, 5h_c; triangle, 5h_oc) and the pairing status by the color (green, paired end; blue, first-in-pair single end read; magenta, second-in-pair single end read). On both pages, results are shown in the four panels: (A) 36-bp reads, (B) 54-bp reads, (C) 72-bp reads, and (D) 101-bp reads.

**Additional File 11. Compendium of figures for *Candida-on\y* data with results separated by read length.** A heatmap with hierarchical clustering with statistical support is shown on page 1 with the condition denoted according to letter code from Table 2, followed by the replicate designation and the pairing status such that (0) paired reads, (1) first-in-read single end read, and (2) second-in-read single end read. A PCA plot is shown on page 2 where the conditions are denoted by the shape (circle, rh2_comp; triangle, rh2_del) and the pairing status by the color (green, paired end; blue, first-in-pair single end read; magenta, second-in-pair single end read). On both pages, results are shown in the four panels: (A) 36-bp reads, (B) 54-bp reads, (C) 72-bp reads, and (D) 101-bp reads.

**Additional File 12. Compendium of figures for *E. coli* data with results separated by read length. A** heatmap with hierarchical clustering with statistical support is shown on page 1 with the condition denoted according to letter code from Table 2, followed by the replicate designation and the pairing status such that (0) paired reads, (1) first-in-read single end read, and (2) second-in-read single end read. A PCA plot is shown on page 2 where the conditions are denoted by the shape (circle, DMEM; triangle, LB) and the pairing status by the color (green, paired end; blue, first-in-pair single end read; magenta, second-in-pair single end read). On both pages, results are shown in the four panels: (A) 36-bp reads, (B) 54-bp reads, (C) 72-bp reads, and (D) 101-bp reads.

**Additional File 13. Compendium of scatterplots for all data sets with results aggregated by read length.** The differentially expressed genes identified using an adjusted p-value (FDR) cutoff ≤0.05 at varying read lengths within a dataset were compared using Pearson’s correlation implemented in the R statistical tool and illustrated as a matrix of scatterplots. The diagonal represents the histogram of log-transformed fold-changes within the comparison. The lower plots represent the correlation between comparisons with singleton DEGs identified for compariosns on the x-axis (pink) and y-axis (green). Genes with FDR > 0.05 in both comparisons are not shown. The upper portion of the plot lists the corresponding Pearson’s correlation coefficient and the number of singleton DEGs identified in each comparison. Each scatterplot is labeled by the comparison according to the letter code from Table 2. A separate plot is shown for paired reads (labelled “0”). first read in pair (labelled “1”), and second read in pair (labelled “2”).

**Additional File 14. Compendium of scatterplots for all data sets with results aggregated by read pairing.** The differentially expressed genes identified using an adjusted p-value (FDR) cutoff ≤0.05 at varying read lengths within a dataset were compared using Pearson’s correlation implemented in the R statistical tool and illustrated as a matrix of scatterplots. The diagonal represents the histogram of log-transformed fold-changes within the comparison. The lower plots represent the correlation between comparisons with singleton DEGs identified for compariosns on the x-axis (pink) and y-axis (green). Genes with FDR > 0.05 in both comparisons are not shown. The upper portion of the plot lists the corresponding Pearson’s correlation coefficient and the number of singleton DEGs identified in each comparison. Each scatterplot is labeled by the comparison according to the letter code from Table 2. A separate plot is shown for the various read lengths.

## MICROBIAL GENOMICS

